# CaliAli, a tool for long-term tracking of neuronal population dynamics in calcium imaging

**DOI:** 10.1101/2023.05.19.540935

**Authors:** Pablo Vergara, Yuteng Wang, Sakthivel Srinivasan, Yoan Cherasse, Toshie Naoi, Yuki Sugaya, Takeshi Sakurai, Masanobu Kano, Masanori Sakaguchi

## Abstract

Neuron-tracking algorithms exhibit suboptimal performance in calcium imaging when the same neurons are not consistently detected, as unmatched features hinder intersession alignment. CaliAli addresses this issue by employing an alignment-before-extraction strategy that incorporates vasculature information to improve the detectability of weak signals and maximize the number of trackable neurons. By excelling in neural remapping and high spatial overlap scenarios, CaliAli paves the way toward further understanding long-term neural network dynamics.

## Main

Existing neuron-tracking algorithms used in one-photon calcium (Ca^2+^) imaging align the spatial footprints of neurons in different recording sessions and apply spatial (CellReg)^1^ and temporal (SCOUT)^2^ similarity thresholds to identify matching neurons. However, active neural populations fluctuate over time^3,4^, which hinders the estimation of brain deformations from unmatching neural features^5^. Indeed, a substantial part of brain misalignment is non-rigid (**Extended Data Fig. 1**), which may cause incorrect alignment of neighboring neurons when neurons are intermittently detectable (**Fig. 1a**). This problem is exaggerated when neurons are tracked over longer periods of time as their footprint projection becomes more dissimilar. To address this issue, we developed CaliAli (<u>Cal</u>cium Imaging intersession <u>Ali</u>gnment), a tool for long-term neural tracking that incorporates information from blood vessels (BVs) and neurons to correct for inter-session misalignment (**Fig. 1b-c, Extended Data Fig. 2**). In artificially misaligned video simulations (**Extended Data Fig. 3**; **Supplementary Video 1, 2**), incorporating shapes of BVs, in contrast to other projections such as neuron shapes, the average frame of the video, or a filtered version of the mean frame, enhanced registration performance and diminished maximum displacement errors (**Fig. 1d-f; Extended Data Fig. 4**). These improvements were observed with BV spatial correlations higher than 0.4 as determined by video simulations with different magnitudes of correlation across sessions (**Fig. 1g-i, Extended Data Fig. 5**). This magnitude of BV correlation was maintained for up to 40 days in Ca^2+^ imaging recordings from mice (**Fig. 1j**)—an interval longer than that used in most long-term neuron-tracking experiments^4^. Even so, if lengthier periods are to be covered, CaliAli can align remote recording sessions using structural information from intermediate recording sessions if inter-session gaps are short (**Supplementary Note 1**). We illustrate this scenario by simulating imaging sessions in which the BV correlation is high between consecutive sessions but low for larger inter-session gaps (**Fig. 1k**) and show that incorporating intermediate recordings between non-alignable recordings markedly improved neuron tracking performance (**Fig. 1l**).

**Fig. 1.**
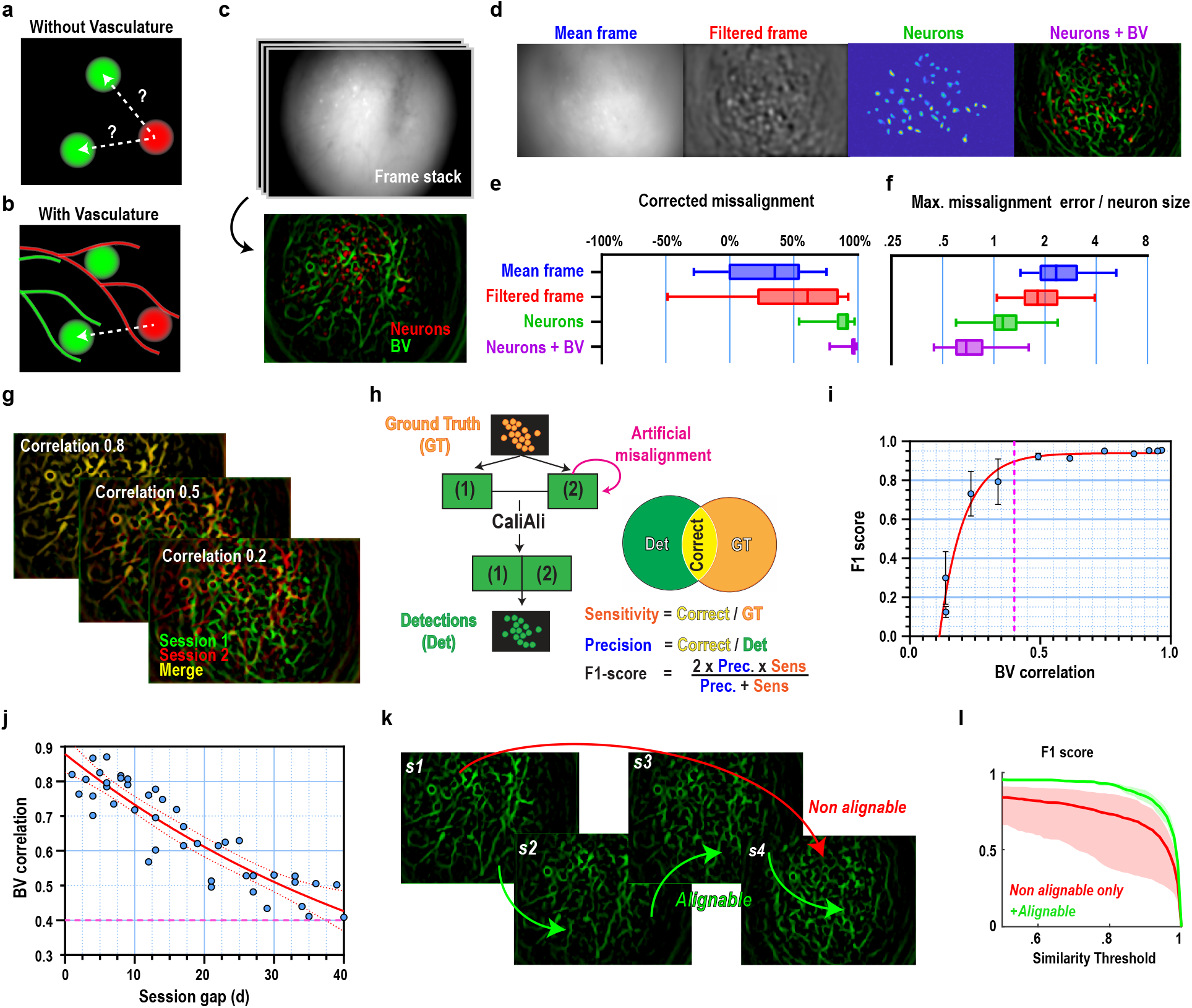
CaliAli incorporates blood vessel (BV) information to maximize neuron tracking performance. **a, b**, Session registration with or without BVs. **c**, BV-neuron projection from a raw frame stack. **d-f**, Comparison of projections and their registration performance. **g**, Simulated BV correlation. **h**, Comparison between ground truth and CaliAli-extracted neurons. **i**, F1 score vs. simulated BV correlation. **j**, BV correlations in recordings from mice vs. inter-session gaps. **I,j**, Red line reflects a sigmoidal fit, and dashed line indicates the threshold where F1 score varies by >5%. **k**, Simulation of BV decorrelation over time. **l**, Tracking performance between the first and last session with or without intermediate sessions. Similarity threshold is the temporal similarity between the extracted component and the matched ground truth neurons used to determine true positives. For **i**, a threshold of 0.8 was used.

A major problem with existing neuron tracking algorithms is their inability to clearly differentiate between inactive and undetected neurons^4^. Unlike footprint alignment methods, CaliAli extracts neural signals from BV-aligned concatenated videos, enabling the identification of Ca^2+^ activity with a low signal-to-noise ratio (SNR) that might otherwise go undetected (**Fig. 2a-c, Extended Data Fig. 6a**). This improves interpretability by ensuring that a consistent number of neurons are tracked across sessions. CaliAli achieves this heightened sensitivity through several optimized modules: preprocessing steps to minimize artifacts at session concatenation points, a rapid initialization pipeline for sparse Ca^2+^ activity, and batch non-negative matrix factorization to reduce memory demands (**Supplementary Note 2, Extended Data Fig. 7**).

**Fig. 2.**
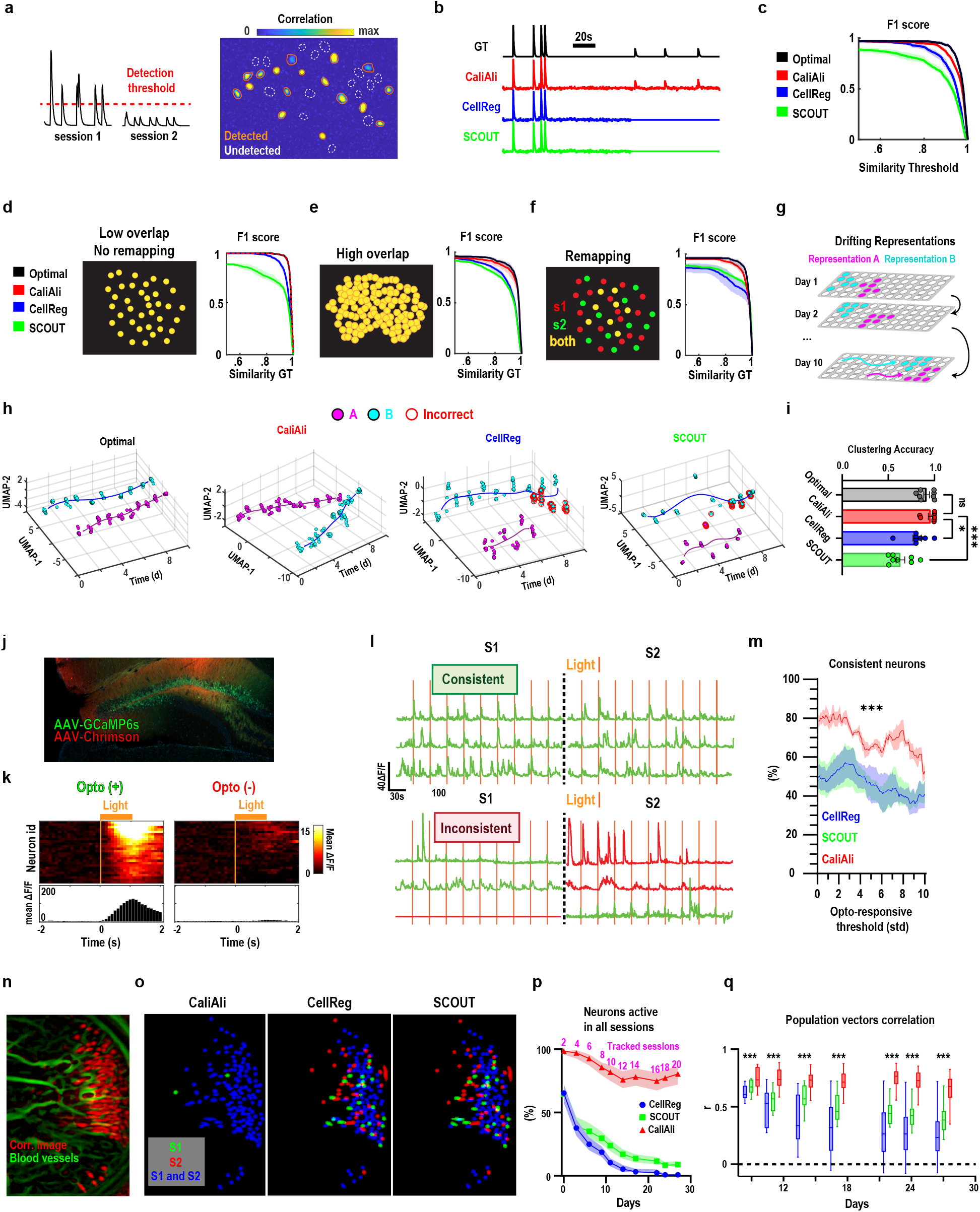
CaliAli maximizes the number of tracked neurons while maintaining consistent properties of tracked neurons. **a**, Simulated scenario in which some neurons’ signal-to-noise ratio fluctuates into undetectable levels. **b,c** Traces for partially undetected neurons and overall tracking performance. **d,e,f**, Tracking performance in different activity scenarios. **a-f** Optimal plot (black) is the maximum achievable performance by CNMFe^10^ (in denoised and perfectly aligned videos). **h**, Dimensionality reduction and unsupervised clustering of Ca^2+^ activity during representational drift. Red circles depict incorrect classifications. **i**, Clustering performance. Repeated measures one-way ANOVA with Dunnet’s multiple comparisons. **j**, Histology of GCaMP6f and Chrismon. **k**, Heatmap and average peristimulus time histograms for opto(+) and opto (-) neurons. **l**, Representative traces for optogenetically consistent and inconsistent neurons. Green = opto(+), red = opto(+). **m**, Percentage of optogenetically consistent neurons for different opto(+) thresholds (for **k**, the threshold is 3 standard deviations). Permutation test. Error bars depict the confidence intervals for different CNMFe initialization parameters applied to data from one mouse. **n**, BV-neuron projection from a 4-week Ca^+2^ imaging experiment. **o**, Spatial components obtained by different methods. **p**, Proportions of tracked neurons across all sessions over time. **q**, Correlation between pairs of population vectors vs. time. Two-way ANOVA with Sidak’s multiple comparisons. \*\*\*\**p* < 0.001, \**p* = 0.018.

We next compared CaliAli’s performance with that of other neural tracking algorithms in three scenarios common to Ca^2+^ imaging experiments: (1) low overlap of neurons with active populations remaining consistent across sessions (**Fig. 2d**), (2) high overlap of neurons (**Fig. 2e**), and (3) remapping of neuronal population activity (**Fig. 2f**). CaliAli performed better than other methods in all scenarios (**Fig. 2d-f, Extended Data Fig. 6b-d**), indicating that it improves trackability in both ideal and challenging conditions. We also simulated a more complex scenario of representational drift (**Fig. 2g**), which is when two orthogonal neural representations gradually change over time while preserving information content^6^. Dimensionality reduction^7^ and unsupervised clustering^8^ of Ca^2+^ traces obtained by CaliAli recapitulated the multi-dimensional structure of the neural trajectories in a more precise manner than other methods (**Fig. 2h, i**). We also tested CaliAli’s performance using actual Ca^2+^ imaging data. We transduced GCaMP6 and the stimulatory opsin Chrimson in dentate gyrus (DG) granule cells (**Fig. 2j**), allowing us to optogenetically tag a subset of Chrimson-positive neurons (49.6% ± 2.8% of neurons) by orange light stimulation (**Fig. 2k**). If neural tracking is accurate, neurons responding to light stimulation in one session would be expected to respond again in a subsequent session (**Fig. 2l**). We found that the proportion of optogenetically consistent neurons was higher with the use of CaliAli versus other methods (**Fig. 2m**), further corroborating its better tracking performance.

Finally, we utilized CaliAli to track DG granule cells when a mouse explored two different contexts over a period of 4 weeks. In contrast to other methods, CaliAli detected more neurons that were active in all sessions (**Fig. 2n-p**). Also, population vectors during different exploration sessions in the same context were more highly correlated for data obtained by CaliAli than by other methods (**Fig 2q**), consistent with observations that the DG produces stable neural representations over time^9^.

In summary, CaliAli is a powerful tool that outperforms state-of-the-art methods in diverse neuron tracking scenarios. It identifies a consistent number of tracked neurons and improves the detectability of smaller Ca^2+^ transients, making it suitable for studying brain dynamics over long periods of time, including representational drift, remote memory processing, neuron development, or behavior measured across multiple trials.

## Materials and Methods

### Animals

All animal experiments were approved by the University of Tsukuba Institutional Animal Care and Use Committee. Mice were kept in a home cage in an insulated chamber maintained at an ambient temperature of 23.5 ± 2.0°C with a 12-h light/dark cycle and ad libitum access to food and water according to institutional guidelines. For simultaneous imaging and optogenetic experiments, we utilized a mouse line (harboring TIGRE-Ins-TRE-loxP-stop-loxP(LSL)-GCaMP6s; Ai94D, stock #024104, Jackson Laboratory, Sacramento, CA, USA). For long-term imaging and video simulation parameterization, virus-injected mice in a C57BL/6J background were used.

### Virus

Adeno-associated viruses (AAVs) were prepared as previously described^11^. For simultaneous imaging and optogenetic experiments, the following viruses were used: AAV1-Syn-Flex-ChrimsonR-Tdtomato (Addgene #62723), AAV2retro-CaMKII-0.4-Cre (Addgene #105558), and AAV2retro-cFos-tTA-pA (Addgene #66794). The AAV2retro-CamKIIa-jGCaMP8s-WRPE virus (Addgene #176752) was used for long-term imaging experiments. The AAV10-CamKII-GCaMP6f-WPRE virus (Addgene #100834) was used to obtain parameters utilized in video simulations and to calculate inter-session missalignment (Extended Data Fig. 1a).

### Virus injection

Mice between 9 and 15 weeks of age were anesthetized using isoflurane and secured in a stereotaxic apparatus (Stoelting, USA). AAV solution (70 nl) was injected into the dorsal hippocampus at AP -2.0 mm, ML +1.2 mm, and DV -1.7 mm relative to bregma. The injection was performed using a Picospritzer III air pressure system (S48 Stumilator, Grass Technologies, USA) connected to a glass pipette injector. The injection process lasted 15 min, after which the injector needle remained in position for 5 min before being gently removed. Following injection, mice were given a minimum recovery period of 1 week before lens implantation.

### Lens implantation

A microendoscope lens (1-mm diameter, 4-mm length, Inscopix, USA) was placed in the dentate (DG) at AP -2.0 mm and ML +1.25 mm relative to bregma and 1.53 mm below the dura. One week after, a UCLA miniscope baseplate^12^ was attached above the implanted lens. After baseplate surgery, mice were habituated to a dummy microendoscope for 1-2 weeks before recording.

### Preparation of tissue sections

After imaging, mice were perfused transcardially with phosphate-buffered saline (PBS; 0.1 M) and 4% paraformaldehyde (PFA). Brains were removed, fixed overnight in PFA, and transferred to PBS. Coronal sections (30 µm) were cut using a vibratome (VT1200S, Leica). Sections were mounted on slides with mounting medium containing DAPI (Merck). Images of GCaMP6s- and ChrimsonR-Tdtomato-expressing neurons were obtained using a Zeiss Axio Observer Z1 microscope.

### Ca^2+^ imaging and optogenetic manipulation

A miniaturized microscope with flexible light source input (Tscope)^13^ was utilized for neuronal imaging and manipulation. For imaging without optogenetic manipulation, we used a blue laser (445 nm, custom-made) delivering 0.3-1.3 mW at the bottom of the Tscope. For the opt-tagging experiment, we used a blue laser (473 nm, Shanghai Laser & Optics Century Co., Ltd., China) delivering 0.07-2.7 mW and an orange laser (589 nm, Shanghai Laser & Optics Century Co., Ltd.) delivering 0.3-1.2 mW at the bottom of the Tscope. Stimulation was delivered through a custom-made laser combiner and an optic patch cable (Thorlabs, Japan). For all experiments, images were acquired at 10 frames/s. Laser intensity, gain, and exposure settings were customized for each mouse while monitoring the fluorescence intensity histogram to ensure that the highest possible dynamic range was achieved without signal saturation. For opto-tagging experiments, the blue laser was used to stimulate GCaMP (300-s sessions), and the orange laser delivered a 1-s pulse every 29 s (10 Hz, 50% duty cycle). We performed two opt-tagging sessions separated by 30 min.

### Long-term imaging

Recordings were made in two distinct environments. Context A consisted of a chamber with white plastic walls and a stainless steel grid (width × depth × height, 310 × 250 × 280 mm). Context B consisted of a circular plastic chamber with a wooden bedding floor (22-cm diameter)^14^. For context A, a white acrylic drop pan under the grid floor was cleaned with 75% ethanol, generating a background odor, whereas no ethanol was employed in context B. Recordings were made every 2-4 days for 28 days. Each day, recordings were performed for contexts A and B, with a 30-min interval between recordings. The order of recordings in A and B was changed each day to maintain a balanced design. Each recording session lasted 5 min.

### Obtaining realistic neural parameters from DG imaging data

Recordings were made 4 days apart from a GCaMP6f-expressing mouse exploring context A for 5 min. Parameters obtained from recorded neurons were used to create video simulations.

### *In vivo* Ca^2+^ imaging data processing

Raw Ca^2+^ imaging videos were spatially downsampled four times. Motion artifacts were corrected utilizing blood vessels (BVs) and the log-demon image registration algorithm. Ca^2+^ traces were extracted by CNMFe using the implementation and preprocessing steps described in **Supplementary Note 2** and **Extended Fig. 8**. The spatial filtering (gSig) was 2.5. For simulations, the minimum peak-to-noise ratio (min_pnr) and minimum pixel local correlation (min_coor) were 2.5 and 0.15, respectively. In the opto-tagging experiment, min_pnr was set to 5 and min_corr varied from 0.4 to 0.8 in increments of 0.05. This was performed to assess the performance of the CaliAli algorithm under different scenarios, ranging from a scenario in which the majority of neurons were extracted (albeit with some false positives) to a scenario in which false positives were minimized at the cost of potentially missing some neurons. For the long-term imaging experiment, it was not feasible to combine different initialization parameters across the 20 recorded sessions due to computational constraints. To address this issue, we manually defined min_pnr and min_corr by carefully monitoring the correlation and PNR image of each recording. To minimize potential bias, we randomly shuffled the session identities during threshold determination. We provide an overlay of the footprints over the correlation image obtained for each recording using the source data included in this paper.

### Statistical analysis

Statistical analysis was performed in MATLAB (MathWorks, Maryland, USA) and Graphpad Prism (GraphPad, California, USA). Error bars depict the 95% confidence interval of the mean across all panels, except for in box-and-whisker plots where they depict the range of values. Shaded error bars were obtained by bias-corrected and accelerated bootstrap. Type I error was set to α = 0.05.

### Video simulations

Spatial components were simulated by randomly sampling from 1,137 DG granule cells from eight GCaMP6f-expressing mice. Spatial components were positioned randomly in the field of view, constrained by minimum distances to neighboring cells (low overlap: 26 µm, medium overlap: 21 µm, high overlap: 8 µm). Temporal components were simulated considering rising times produced by a Bernoulli process and subsequently convolved with a temporal kernel *g* (*t*) = exp(− *t*/τ_*d*_) − exp(− *t*/τ_*r*_). The Ca^2+^ rates and kinetics for each neuron were sampled from a lognormal distribution with parameters obtained from mouse DG recordings: transient probability µ = -4.9, σ = 2.25; τ r^-1^ µ = 2.08, σ = 0.29; τ d^-1^, µ = 0.55, σ = 0.44. Note that the mean transient rate in our simulation was marginally higher than that in the empirical data, as neurons were required to exhibit a minimum of one Ca^2+^ transient per session. A constant peak-to-noise ratio of 2 was used in all simulations. Local background fluctuations were modeled using a 2D Gaussian-filtered version of the spatial components (σ = 20), with weakly correlated noise generated by applying a 2D Gaussian filter with σ = 0.5 on white noise. Inter-session misalignment was emulated using gradients of a random 2D Gaussian (σ = 60), scaled to produce a maximum non-rigid misalignment of 15-25 µm, which corresponds to our estimation in the emperical data (Extended data Fig. 1).

Remapping was simulated by rendering a subset of neurons inactive in certain sessions. Variable signal-to-noise ratios were simulated in a similar manner as in remapping, but the amplitude of Ca^2+^ transients was reduced by 80% instead of inactivating neurons. The parameters employed in each simulation are found in the source data accompanying this paper.

### Incorporation of realistic BV structures

We utilize frames obtained from DG recordings as a static baseline. To incorporate modest variation in the static baseline, we utilized frames obtained 4 days apart, which were manually aligned and used as a static baseline for each session. For simulations utilizing more than two sessions, additional baseline frames were created by linear interpolation. To simulate BV variation larger than that observed within 4 days (Fig. 1i,k), we computed weighted averages between baseline images obtained from different mice.

### Enhancement of BV structures

BV structures were enhanced from raw one-photon Ca^+2^ imaging frames using Hessian-based enhancement filters. In Ca^+2^ imaging experiments, BVs typically appear as elongated structures with lower intensity values than the surrounding tissue. Hessian-based enhancement filters exploit these characteristics by examining the eigenvalues of the Hessian matrices of an image. The Hessian matrix has two eigenvalues at each pixel location. The relationship between these eigenvalues helps identify different structures in the image. In the case of BVs, the primary eigenvalue (λ1) is generally much smaller in magnitude than the secondary eigenvalue (λ2), indicating a tubular structure. In practice, the filtering is implemented as follows:

Step 1: Empirically determine the range of BV diameters d_1_, d_2_, …d_n_ in the raw image.

Step 2: For each diameter, perform steps 3 to step 6.

Step 3: Sequentially convolute the columns and rows of the image using a 1-D Gaussian filter with 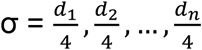

Step 4: Compute the Hessian matrix of the filtered image:

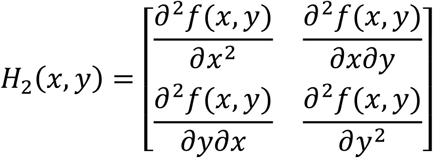

Here,*f* (*x,y*) is the intensity function of the filtered image at pixel (*x,y*) and, 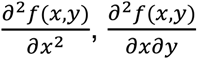 etc. are second-order partial derivatives of *f*. For faster computation, we calculated the Hessian matrix using the implementation described by Yang et al.^15^

Step 5: Find the eigenvalues of the Hessian matrix. The eigenvalues of *H*_2_(*x,y*) are obtained from the following analytical equation:

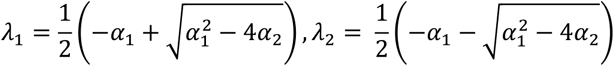

Here, *α*_1_ and *α*_2_ are the roots of the characteristic polynomial of given by:

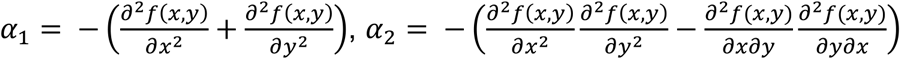

Step 6: Calculate the filter response function, defined as the larger absolute value between λ_1_ and λ_2_ :

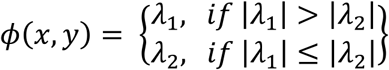

Step 7: To obtain the enhanced vasculature image, we applied the filter response function to the original image by combining the results from multiple scales (i.e., different vessel diameters or filter sizes):

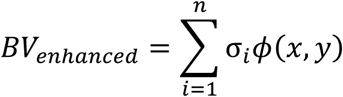

We considered 10 diameter sizes ranging from 2.4×gSiz to 3.5×gSiz, where gSiz is the filter size (in pixels) defined in CNMFe.

### Neuron tracking parameters

We utilized default parameters for SCOUT and CellReg except for a non-rigid registration approach for field of view alignment, as all simulations involved non-rigid deformations. In some cases, the footprint registration algorithm used by SCOUT produced lower performance than CellReg, mainly when neural densities were low and the active neural population remapped. To ensure that any differences in performance were not due to incorrect footprint alignment, we used the same alignment module as was used with CellReg for SCOUT. Through these modifications, we were able to maintain or improve the performance of SCOUT.

### Dimensionality reduction and unsupervised clustering of drifting neural activities

The population activities shown in Fig. 2h,i were subjected to dimensionality reduction and clustering using the UMAP algorithm. Cosine distance metric, a minimum distance of 0.1, and 10 nearest neighbors were used for this purpose, as they have been shown to produce the best outcomes in ground truth data. Unsupervised clustering was performed on the reduced data using the k-means algorithm and squared Euclidean distance metric.

### Inter-session registration

Sessions were registered using the diffeomorphic log-Demons algorithm^16^, which is an image registration method that computes a smooth and invertible deformation field to match a moving image to a fixed reference image. The algorithm is based on the original Demons algorithm by Thirion^17^, which was extended to ensure diffeomorphism (i.e., smoothness of the deformation). The optimal displacement field *S* that aligns a moving image M to the static image *F* is estimated by minimizing the velocity field 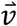 in the following energy function:

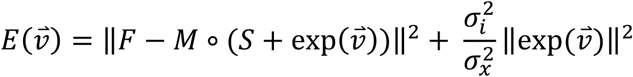

The velocity field 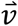 is used to additively update *S* in each iteration, and 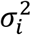 and 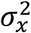 are weights regulating the similarity term 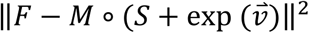 and the maximum step in each iteration, respectively. The algorithm uses an exponential map to update the deformation field to ensure that it remains diffeomorphic. Here, 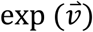 is the exponential map of the vector field 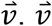. is calculated using the additive demon algorithm: 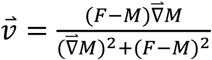

Two regularization steps are included at each iteration. For a fluid-like regularization, we convolute 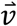with a Gaussian kernel with size =*σ*_*fluid*_. For a diffusion-like regularization, we convolute *S* with a Gaussian kernel with size = *σ*_*diffusion*._ The regulation parameters used at each registration level are 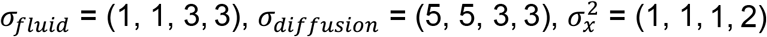 and 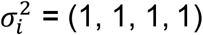

## Code availability

The codes for CaliAli, along with demo videos and tutorials, are available on GitHub: https://github.com/CaliAli-PV/CaliAli

## Source data

The Ca^2+^ imaging videos supporng this study and the codes to recreate simulaons are available at: hps://data.mendeley.com/datasets/mvg4w89s4s/dra?a=febfa476-d9ef-4e98-abe4-24113a24549e

## Acknowledgments

We thank K.G. Akers for comments on the manuscript. We thank M. Sakurai for secretarial support.

