## Supplementary figures and notes for "CaliAli, a tool for long-term tracking of neuronal population dynamics in calcium imaging"

### Figure E1

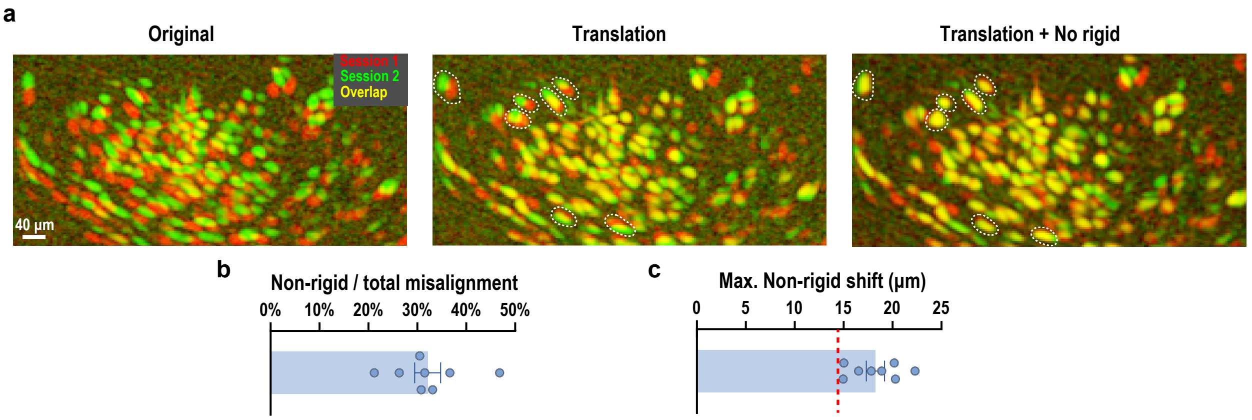


**Extended Data Fig 1 | Non-rigid misalignment in dentate gyrus recordings. a**, Correlation image of two sessions (4-day inter-session gap) before alignment, after translation, and after translation and non-rigid registration. White dashed lines show neurons incorrectly aligned by translation but not by non-rigid registration. **b**, 30% of the total misalignment was non-rigid. **c**, Maximum non-rigid misalignment. Red line indicates the average size of a granule neuron. Across all mice, non-rigid misalignments were larger than the average granule neuron, which would result in incorrect neuron tracking. n = 8 mice.

#### Figure E2

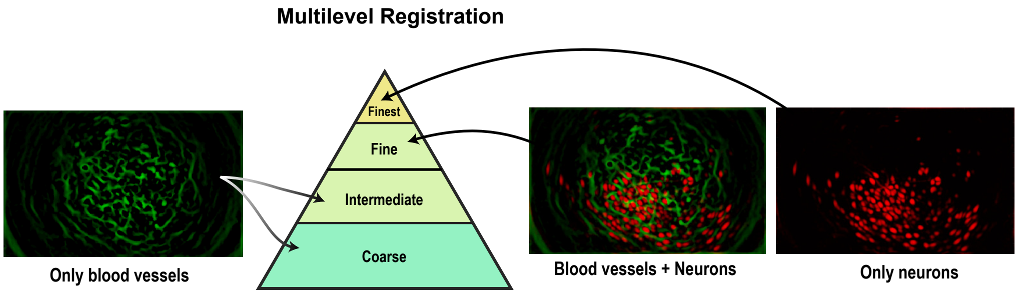


**Extended Data Fig 2 | Multilevel registration utilizing blood vessel (BV) and neuron shapes.** Inter-session alignment performed with log-demon registration. This algorithm calculates a spatial transformation obtained from image gradients that maximizes similarities between two images. In the presence of large displacement–and given the homogenous shapes of neurons–this approach is susceptible to local minima (e.g., the algorithm stops when two non-corresponding neurons partially overlap). To address this, we utilized a multilevel approach in which larger displacements are calculated on coarser versions of BV images, and precise adjustments are calculated using neuron shapes. In contrast to neurons, BVs display higher structural heterogeneity and are less susceptible to local minima; therefore, they are more suitable for correcting larger displacements in the field of view.

#### Figure E3

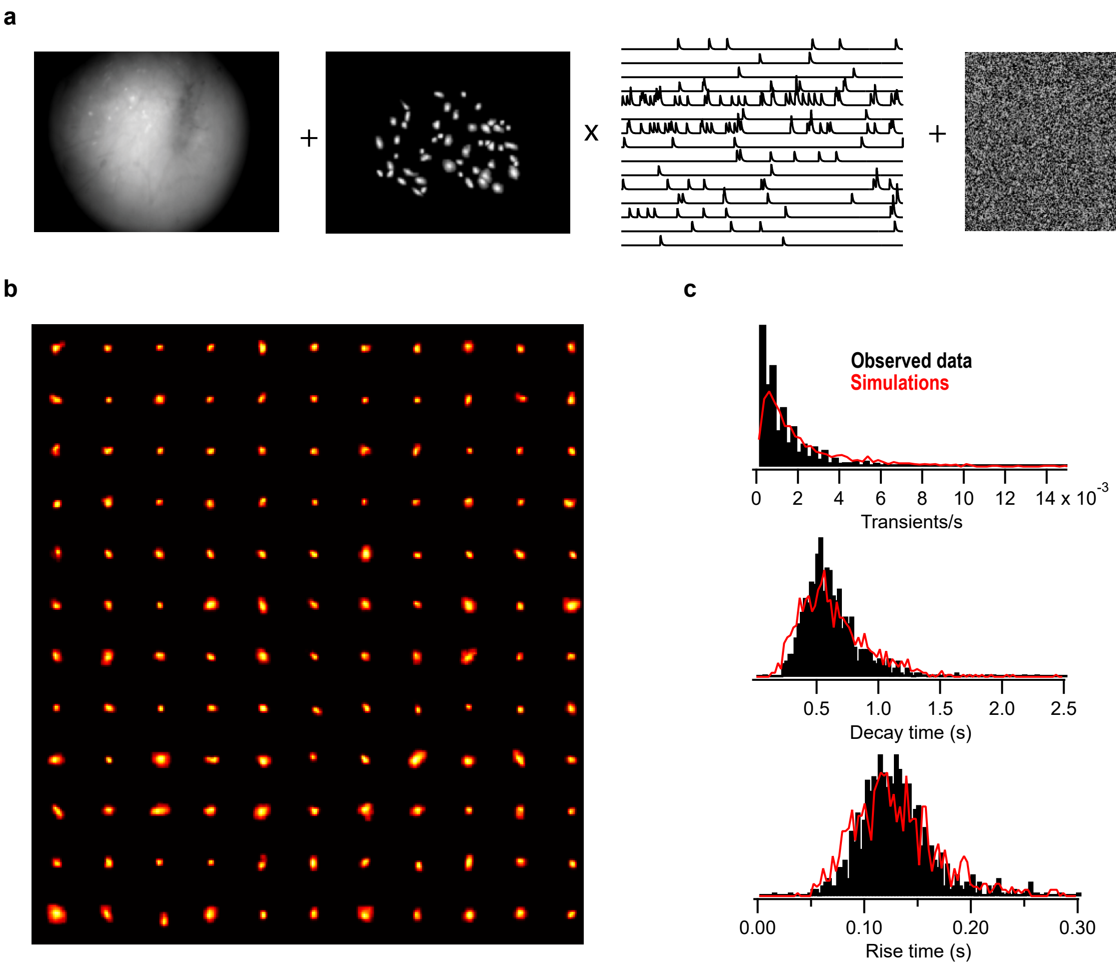


**Extended Data Fig 3 | Video simulations**. **a**, Video simulations combined a constant baseline, neuron spatial and temporal component, noise, and local background fluctuations. **b**, Spatial components were simulated by randomly sampling from a collection of 1,137 dentate gyrus granule cells obtained across eight mice. **c**, Neuron mean transient rates and $Ca^{2+}$ decay and rise times were randomly selected from a lognormal distribution with parameters estimated from actual granule neuron data.

#### Figure E4

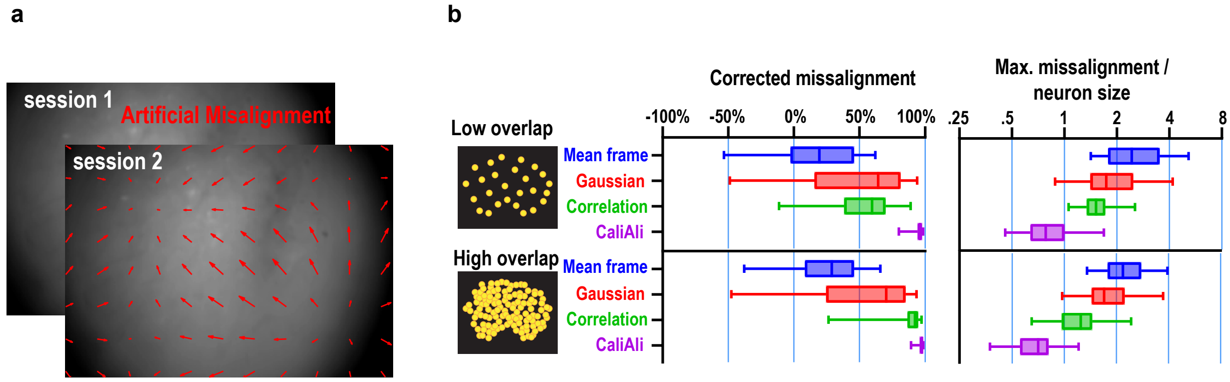


**Extended Data Fig 4 | Inter-session alignment performance using different projections**. **a**, Artificial inter-session misalignments were introduced in one of the two video sessions. **b**, Registration performance of different projections under different neural densities. Moderate neural densities are shown in Fig. 1e.

#### Figure E5

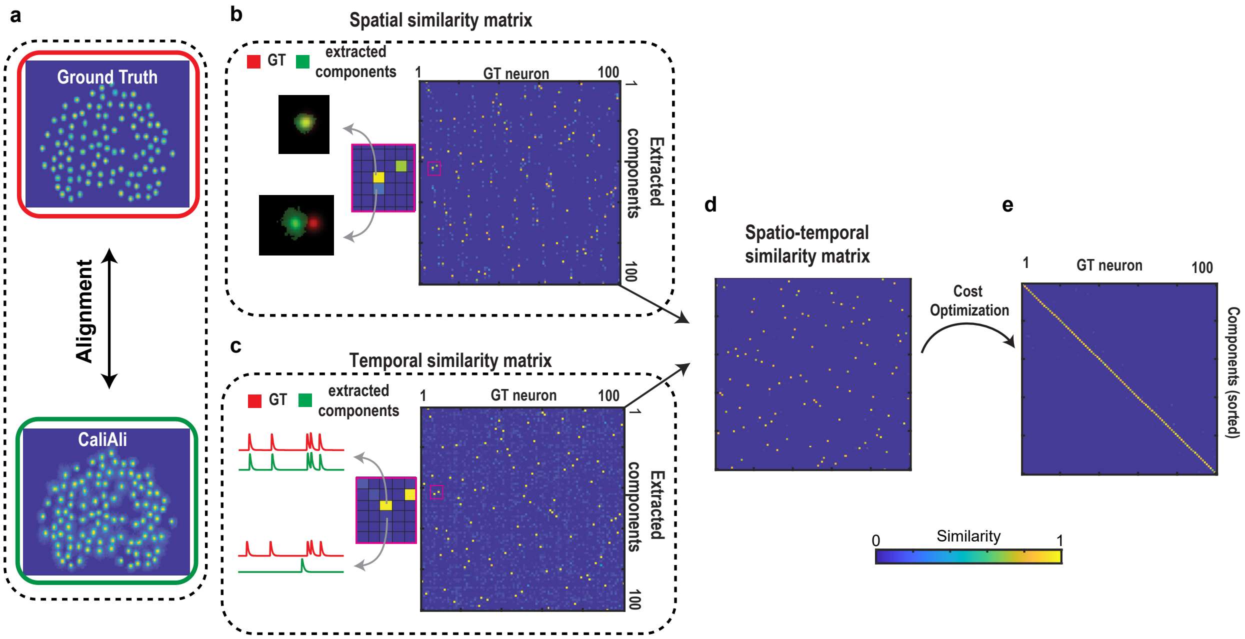


**Extended Data Fig 5 | Matching of extracted components with ground truth (GT) neurons.** GT and extracted components were matched as follows: **a**, A projection of the extracted component was aligned with the GT footprints. **b**, A spatial similarity matrix (i.e., cosine similarity) for each combination of GT and extracted neuron was calculated. **c**, The same was done for temporal components. **d**, An element-wise product of both matrices was calculated to create a spatiotemporal matrix. **e**, The rows and columns of the spatiotemporal matrix were permutated to find the assignment maximizing the similarity between GT and extracted components. This linear assignment problem was solved using the matchpair function in MATLAB. This approach ensures one-to-one matching and is suitable for cases in which the number of extracted components differs from the number of GT neurons (i.e., non-matched components have zero spatiotemporal similarity). Finally, we used the temporal similarity of the matched components as a measure of tracking performance. Matched components with temporal similarity above 0.8 were considered true-positive.

#### Figure E6

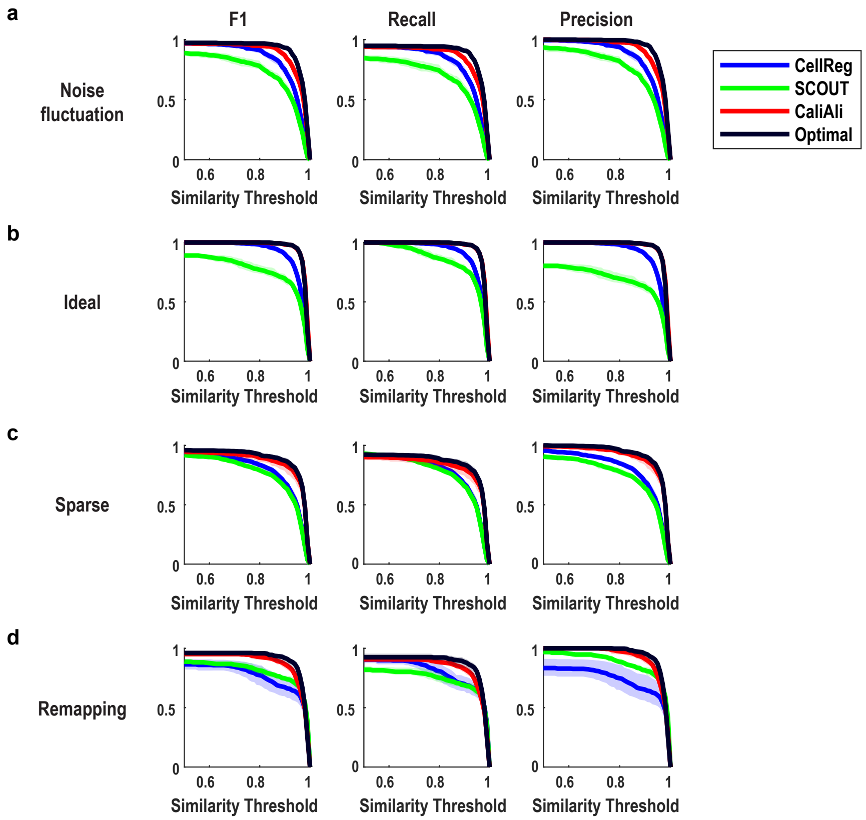


**Extended Data Fig 6| Ground truth performance scores.** F1, recall, and precision scores for Fig. 2c-g.

### Figure E7

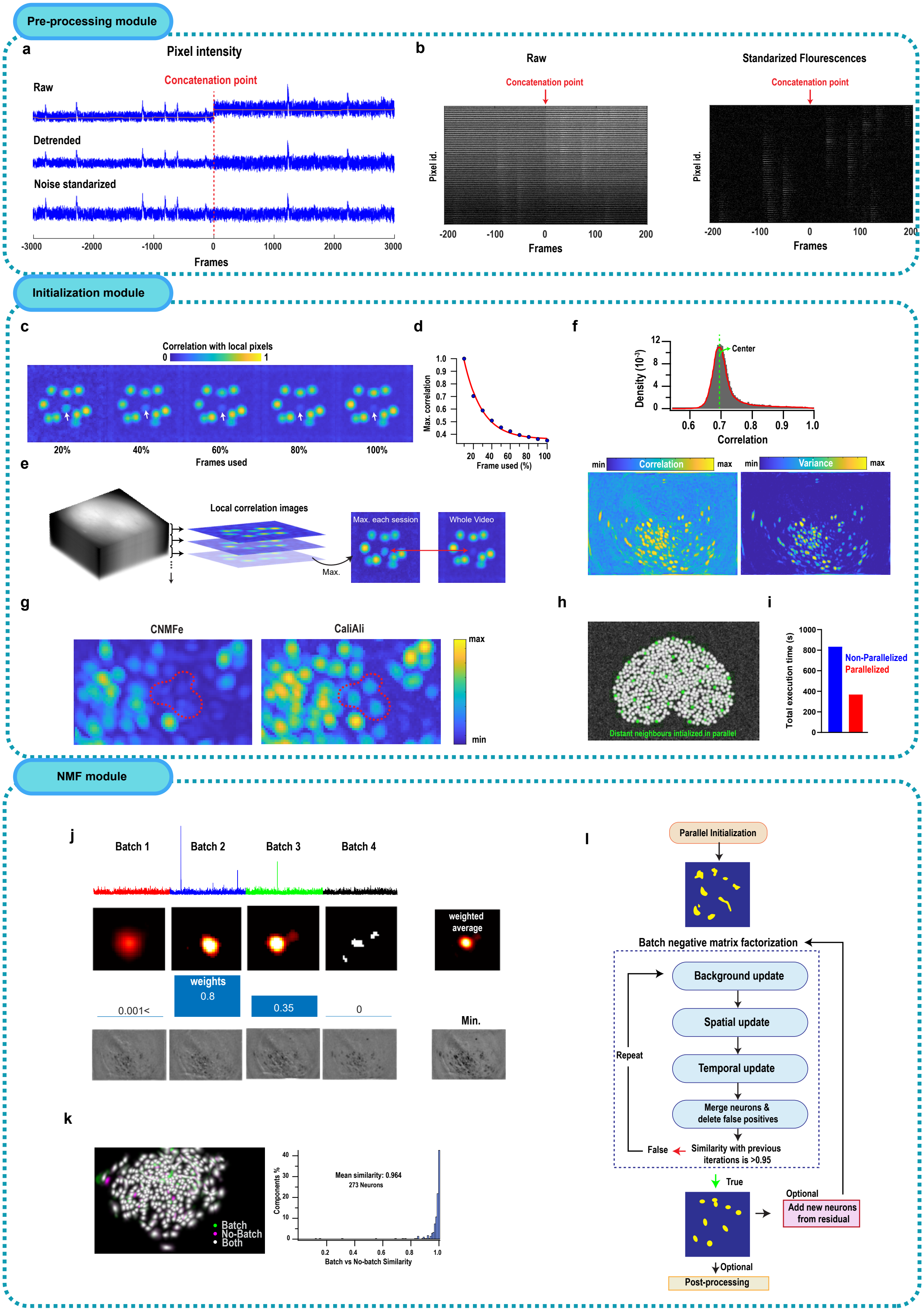

Pre-processing module
a
Pixel intensity
Concatenation point
Raw
Detrended
Noise standarized
-3000
-2000
-1000
0
1000
2000
3000
Frames
b
Raw
Concatenation point
Pixel id.
-200
-100
0
100
200
Frames
Standarized Flourescences
Concatenation point
Pixel id.
Frames
Initialization module
c
Correlation with local pixels
0
1
20%
40%
60%
80%
100%
Frames used
d
Max. correlation
Frame used (%)
e
Local correlation images
Max. each session
Whole Video
Max.
f
Density (10⁻³)
Center
Correlation
Correlation
Variance
min
max
g
CNMFe
CaliAli
max
min
h
Distant neighbours intialized in parallel
i
Total execution time (s)
Non-Parallelized
Parallelized
NMF module
j
Batch 1
Batch 2
Batch 3
Batch 4
weighted average
weights
0.8
0.001<
0.35
0
Min.
k
Batch
No-Batch
Both
Mean similarity: 0.964
273 Neurons
Components %
Batch vs No-batch Similarity
l
Parallel Initialization
Batch negative matrix factorization
Background update
Spatial update
Temporal update
Merge neurons & delete false positives
Repeat
False
Similarity with previous iterations is >0.95
True
Optional
Add new neurons from residual
Optional
Post-processing

**Extended Data Fig 7 | Preprocessing, initialization, and non-negative matrix factorization modules utilized by CaliAli. a**, Representative pixel intensity trace for the raw signal, after detrending, and after noise scaling**. b**, Vectorized pixel intensities around the concatenation point for the raw signal and after fluorescence standardization. **c**, Correlation image calculated from different fractions of the total video length when a neuron (white arrow) is only transiently active. **d**, Maximum signal in the correlation image for the neuron shown in **c**. **e**, CaliAli computes a correlation image by processing a smaller number of frames at a time (i.e., batch) and then generates a maximum projection for each. **f**, To minimize the effect of spurious correlation, the correlation image is centered at zero and subsequently squared. **g**, Correlation images obtained by CaliAli and standard CNMFe from concatenated dentate gyrus recordings. **h**, Distant neighbor initialized in parallel. **i**, Execution times for non-parallelized vs. parallelized initialization (50,000 frames). **j,** Summarization of temporal, spatial, and background components obtained in batch mode. **k,** Components obtained with and without batch implementation. **l**, Overall pipeline utilized by CaliAli.

**Supplementary note 1 | Multisession registration algorithm.** To align multiple $Ca^{2+}$ imaging sessions, each session must be transformed into a common coordinate system. Previous methods for neuron-tracking algorithms, such as CellReg[1] and SCOUT[2], employ two approaches: aligning each session to a common reference session or sequentially aligning each session with its predecessor. However, these approaches have limitations; slow but consistent changes in the field of view make it difficult to select a suitable reference for each session, and sequential alignment is susceptible to error propagation, as each transformation depends on the previous one. To overcome these limitations, we implemented an iterative image registration approach[3], where multiple images are aligned by calculating all possible pairwise transformations between them, after which a common reference space or atlas is found that minimizes differences among all images. This approach iteratively weights transformations based on the post-registration similarity of images. The alignment of session *n* to the reference space is given by:

$$T_{n \to RS} = \frac{1}{N} \sum_{i=1}^{N} T_{n \to i} \cdot W_{\mathrm{n}}$$

Here, $T_{n \to i}$ is the displacement field that aligns session *n* with session $i$. $W_{\mathrm{n}}$ is a weight matrix given by $W_{\mathrm{n}} = w_{\mathrm{n}}^{l} \cdot w_{\mathrm{n}}^{g}$, in which $w_{\mathrm{n}}^{l}$ is the post-registration average local blood vessel correlation within a radius of 25 pixels of session *n* with each other session, and $w_{\mathrm{n}}^{g}$ is the average global correlation. In practice, we first perform an unweighted registration to determine $W_{\mathrm{n}}$.

This approach can be particularly useful in datasets with high variability or noise, in which not all image pairs may be equally informative for registration.

**Supplementary note 2 | CaliAli is optimized to extract neural signals from long concatenated $Ca^{2+}$ imaging videos.**

Detrending and noise scaling: In existing neuron extraction packages, such as CNMFe, basic functionalities are provided to process concatenated video files. However, artifacts arising from variations in baseline fluorescence across sessions can hinder subsequent neural extraction. To mitigate this issue, we initially applied detrending and noise-scaling to each session (**Extended data Fig. 7a**), effectively reducing artifacts at concatenation points between sessions (**Extended data Fig. 7b**). In scenarios in which neuron and background separation proves particularly challenging, such as when optogenetics and imaging are combined, we additionally employed the robust background subtraction and denoising module featured in the $Ca^{2+}$ imaging pipeline MIN1PIPE[4].

Initialization of transiently active neurons: A prerequisite for CNMFe is an initial estimation of neuron locations, which involves detecting peaks in the local correlation image. However, in one-photon imaging, neuron pixels only correlate when a neuron is active[5]. Consequently, when a neuron is active in just a few sessions, video concatenation often leads to a reduced signal-to-noise ratio (SNR) in the correlation image. We illustrate this case by considering a concatenated recording of 50,000 frames in which a neuron was active in only the first 20% of frames (**Extended data Fig. 7c**). Although this neuron can be effectively extracted if only the first 20% of frames are processed, the more frames are processed, the worse its SNR in the correlation image (**Extended data Fig. 7d**). CaliAli improves the original CNMFe algorithm by sequentially calculating a correlation image from shorter video segments and then calculating a maximum projection (**Extended data Fig. 7e**). As the SNR of the correlation image may vary across sessions, each session's correlation image is centered at zero by subtracting a noise estimate defined as the largest peak below the median of the signal (**Extended data Fig. 7f**). The correlation images are then transformed into variance images (squared correlation), which minimizes variation in background noise levels across correlation images (**Extended data Fig. 7g**).This approach enhances the detection of neurons in extended dentate gyrus $Ca^{2+}$ imaging recordings, providing CaliAli with a distinct advantage over other methods for processing long concatenated recordings.

Parallelized initialization: Initialization is one the most time-consuming steps in the neural extraction pipeline of CNMFe. The original CNMFe implementation initializes neurons sequentially from high-to-low SNRs. Once a neuron is initialized, a rough estimate of this neuron signal is subtracted from the video, and the correlation image is locally recalculated. This improves the initialization performance of dim neurons masked by bright signals. However, this sequential approach is not parallelized and is time-consuming. To expedite initialization, we implemented a module in which neurons are initialized in parallel by choosing distant neighbors (**Extended data Fig. 7h**). This approach is based on the idea that bright neurons only influence the initialization of near neighbors. This reduces execution times to less than half while locally preserving a high-to-low SNR sequential initialization (**Extended data Fig. 7i**).

Batch non-negative matrix factorization (NMF):

CaliAli employs the NMF implementation delineated in CNMFe, as detailed in the original paper[6]. Succinctly, this implementation leverages NMF to iteratively update spatial, temporal, and background components of $Ca^{2+}$ imaging videos. We adapted this implementation to facilitate a batch mode, consequently reducing the memory required for processing concatenated video sequences. Although the original CNMFe incorporates batch implementation, it differs from our approach in that the entire extraction pipeline, including initialization, is performed in batch. This presents the disadvantage that the initialization of a new component requires recalculating NMF iterations for all other batches, causing longer computation times and a heightened risk of error propagation. In fact, the batch implementation of CNMFe frequently leads to suboptimal tracking performance across sessions[2].

By contrast, CaliAli carries out global initialization, enabling batch processing without the need to initialize neurons in each batch. For updating temporal, spatial, or background components, an operation is executed to summarize results across each batch (**Extended data Fig. 7j**): temporal components are concatenated; spatial components are averaged, weighted by the mean temporal activity in each batch; and background parameters are averaged except for the constant baseline, which is expressed as the minimum across all sessions. Our batch implementation's components yielded an average similarity of 0.95 with those obtained by processing the entire video, demonstrating that our approach negligibly affects results while reducing memory requirements (**Extended data Fig. 7k**). Additionally, we developed a straightforward method for determining the optimal number of NMF iterations; after each iteration, updated spatial and temporal components are compared with those from previous iterations (**Extended data Fig. 7l**). If similarity surpasses 0.95, we considered the model to have converged and ceased updating.
